# Group II truncated haemoglobin YjbI prevents reactive oxygen species-induced protein aggregation in *Bacillus subtilis*

**DOI:** 10.1101/2021.05.28.446166

**Authors:** Takeshi Imai, Ryuta Tobe, Koji Honda, Mai Tanaka, Jun Kawamoto, Hisaaki Mihara

## Abstract

Oxidative stress–mediated formation of protein hydroperoxides can induce irreversible fragmentation of the peptide backbone and accumulation of cross-linked protein aggregates, leading to cellular toxicity, dysfunction, and death. However, how bacteria protect themselves from damages caused by protein hydroperoxidisation is unknown. Here we show that YjbI, a group II truncated haemoglobin from *Bacillus subtilis*, prevents oxidative aggregation of cell-surface proteins by its biologically unprecedented protein hydroperoxide peroxidase-like activity, which removes hydroperoxide groups from oxidised proteins. Disruption of the *yjbI* gene in *B. subtilis* lowered biofilm water repellence and the mechanical stiffness of the cell surface, which associated with the cross-linked aggregation of the biofilm matrix protein TasA. YjbI was localised to the cell surface, and the sensitivity of planktonically grown cells to generators of reactive oxygen species was significantly increased upon *yjbI* disruption, suggesting that YjbI pleiotropically protects labile cell-surface proteins from oxidative damage. YjbI removed hydroperoxide residues from a model oxidised protein substrate, bovine serum albumin, and prevented its oxidative aggregation *in vitro*. These findings provide new insights into the role of truncated haemoglobin and the importance of hydroperoxide removal from proteins in the survival of aerobic bacteria.

## Introduction

Truncated haemoglobins (trHbs) are small haem proteins found in microbes and plants (Wittenberg et al., 2002, Boechi et al., 2010). They belong to the globin superfamily and form a distinct family separated from bacterial flavohemoglobins, *Vitreoscilla* haemoglobin, plant symbiotic and non-symbiotic haemoglobins, and animal haemoglobins (Vuletich and Lecomte, 2006, Boechi et al., 2010). The haem group enables trHbs to bind to small biomolecules, such as oxygen (O_2_), carbon monoxide (CO), and nitric oxide (NO) (Wittenberg et al., 2002). Additionally, haem-pocket residues are highly structurally variable, conferring trHbs with diverse roles, such as scavengers (Ouellet et al., 2002) or transport carriers (Liu et al., 2004, Wittenberg et al., 2002) of the corresponding small ligands. Such versatility is possibly related to defence mechanisms against the harmful effects of reactive oxygen species (ROS). trHbs are also implicated in the pathogenicity of some bacteria (Pawaria et al., 2007, Ascenzi et al., 2008, Wittenberg et al., 2002), but the molecular functions of these proteins are elusive.

The trHb family proteins are further divided into three subfamilies, Groups I (trHbN), II (trHbO), and III (trHbP), with each subfamily having different structural characteristics (Bustamante et al., 2016, Wittenberg et al., 2002). Among these subfamilies, trHbO proteins evolutionarily originated before trHbN and trHbP proteins and are distributed into Actinobacteria, Proteobacteria, Firmicutes, and plants (Vuletich and Lecomte, 2006, Boechi et al., 2010). A trHbO from *Mycobacterium tuberculosis* exhibits inefficient gas-exchange capacity with a very slow release of O_2_ (Ouellet et al., 2003) and a slower reaction rate with NO than other trHb subfamilies (Ouellet et al., 2003), suggesting that trHbOs play additional roles other than transporting these molecules (Giangiacomo et al., 2005). In accordance, trHbOs from *M. tuberculosis* (Ouellet et al., 2007), *Thermobifida fusca* (Torge et al., 2009), and *Roseobacter denitrificans* (Wang et al., 2015) can exhibit peroxidase activity on hydrogen peroxide *in vitro.* However, because these bacteria express catalases that scavenge hydrogen peroxide (Manca et al., 1999), the physiological relevance of the peroxidase activity of trHbOs remains unclear.

The gram-positive bacterium *Bacillus subtilis* possesses a trHbO, termed YjbI. YjbI has been heterologously expressed as a recombinant protein in *Escherichia coli,* and the purified protein has been characterised (Choudhary et al., 2005). It exists as a monomer with a molecular mass of 15.1 kDa and comprises 132 amino acid residues (Choudhary et al., 2005). The three-dimensional crystal structure of YjbI in the form of the cyano-Met derivative was determined at 2.15-Å resolution, revealing intriguing structural features, including a shallow depression on the proximal side of the haem. The binding properties of YjbI to O_2_ (Boechi et al., 2010, Ouellet et al., 2002), CO (Boechi et al., 2010, Ouellet et al., 2002, Choudhary et al., 2005, Feis et al., 2008, Lapini et al., 2012), and hydrogen sulphide (Nicoletti et al., 2010) have been extensively studied *in vitro*. Some have observed that YjbI has a high affinity for O_2_ and hydrogen sulphide, and the oxygenated derivative is significantly stable, which may rule out a role of YjbI in O_2_ transport/storage. As is common for many haem proteins, an inherent low peroxidase activity of YjbI on hydrogen peroxide has been reported (Choudhary et al., 2005), but this observation remains controversial. The *yjbI* gene is part of the *yjbIH* operon. YjbH is a bacterial adaptor protein required for efficient degradation of the disulphide-stress–activated transcription regulator Spx by the protease ClpXP (Larsson et al., 2007, Awad et al., 2019). Although these previous studies have implied a possible link between YjbI and response to oxidative stress, the exact function of YjbI remains elusive.

In this study, we present *in vivo* and *in vitro* analyses of the physiological function of *B. subtilis* YjbI. We show *yjbI* is required for the formation of biofilms with normal stiffness, wrinkles, and water repellence. YjbI was found to suppress the ROS-mediated aggregation of the major biofilm matrix protein TasA and localise to the bacterial cell surface. We show that, besides a role in biofilm maintenance, YjbI functions as an antioxidant protein. Moreover, we observed YjbI exerted a peroxidase-like activity on a protein hydroperoxide substrate. YjbI is proposed to function as an antioxidant protein in protecting cell-surface proteins from ROS-induced aggregation via its protein hydroperoxide peroxidase-like activity.

## Results

### *yjbI* disruption reduces the mechanical strength of *B. subtilis* biofilm

When the *yjbI* disruptant (BFS2846, Supplementary Table 1) of *B. subtilis* 168 was cultured in a biofilm-promoting minimal (MSgg) liquid medium, we found that the pellicles, which are floating biofilms formed at the air-liquid interface, of the mutant were more fragile than those of the wild-type (WT) strain, and they were easily dispersed by scooping with a spatula (data not shown). Using atomic force microscopy (AFM), we measured the stiffness of the cell surfaces of the WT and *yjbI* disruptant. The cell surface of the WT bacteria from pellicles was stiffer than the surrounding matrix, as indicated in the AFM phase images by the relatively lower brightness of the cell surface (Fig. 1a). Conversely, the cell surface of the *yjbI* disruptant pellicles showed lower stiffness than the matrix (Fig. 1b), suggesting that the mutant pellicles have a less stiff cell surface than the WT. In addition, the average size of the *yjbI* disruptant cells (4.4 ± 0.13 μm) was significantly greater than the WT cells (3.1 ± 0.16 μm) *(p* < 0.001, mean ± SEM, n = 50) (Fig. 1a and 1b). Introducing a plasmid encoding *yjbI* into the *yjbI* disruptant partially rescued the effect of *yjbI* disruption, resulting in cell-surface stiffness higher than the stiffness of the matrix and an average length of 3.8 ± 0.19 μm (Fig. 1c). These results suggest *yjbI* is important for maintaining the cell-surface stiffness of pellicles. The decrease in the stiffness of the cell surface upon the *yjbI* disruption presumably elongated the cells.

**Fig. 1.**
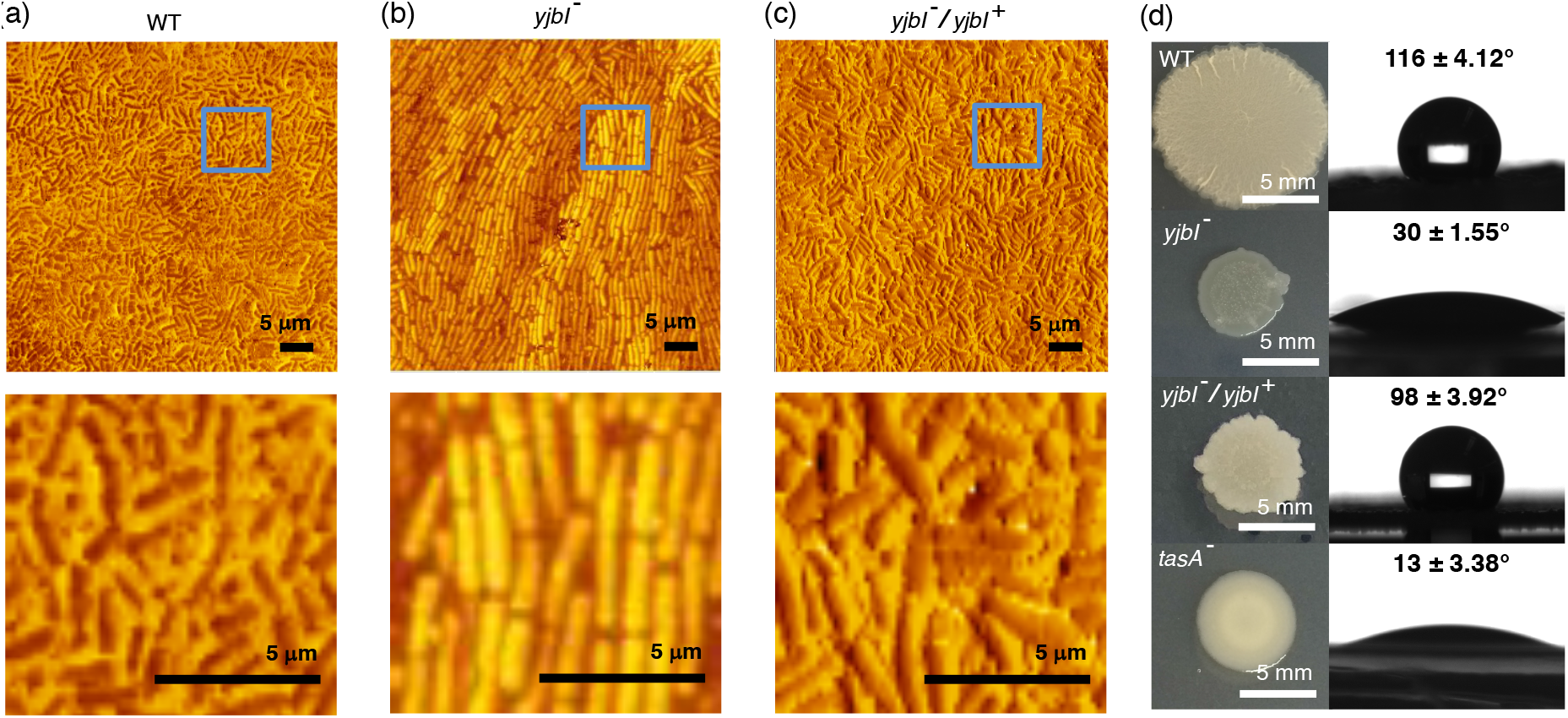
YjbI is needed for normal biofilm formation. Phase-imaging atomic fnrce Smicroscopy (AFM) was used to analyse the wild-type (WT) (a), *yjbI*-disrupted mutant (*yjbI^-^*) (b), and *yjbI^-^* complemented with a *yjbI*-carrying plasmid *(yjbI^-^/yjbI^+^*) (c) of *B. subtilis* biofilm cells from the pellicles on liquid MSgg medium. The areas indicated with cyan boxes in the upper panels were magnified to produce the images in the lower panels in (a), (b), and (c). The contrast represents the relative stiffness of the objects; the objects with dark pixels are stiffer than those with bright pixels within the same image. Similar AFM results were obtained from two independent experiments. (d) Images showing the morphology (left panels) and surface repellency (right panels) of the biofilms of the WT, *yjbI^-^, yjbI^-^/yjbI^+^,* and *tasA*-disrupted mutant (*tasA^-^*) strains of *B. subtilis.* The water contact angles indicated in the right panels represent the mean ± SD of three independent experiments (n = 3).

### *yjbI* is required for normal formation of the *B. subtilis* biofilm

We also observed unusual biofilm formation by the *yjbI* disruptant on a solid medium. Specifically, the *B. subtilis* 168 WT strain formed architecturally complex colonies with characteristic wrinkles on the MSgg solid medium, whereas the *yjbI* disruptant failed to establish such biofilms; instead, it formed flat and glossy colonies (Fig. 1d). Because of the water-repellent properties of the mature biofilm surfaces, the functional integrity of a biofilm can be inferred from the contact angle of a water droplet placed on top of the biofilm (Arnaouteli et al., 2017, Arnaouteli et al., 2016, Kobayashi and Iwano, 2012). Through this method, we examined the integrity of the biofilms of the *yjbI* disruptant. Water droplets placed on the surfaces of the WT biofilms assumed nearly spherical shapes with a contact angle of 116 ± 4.12°, indicating the remarkable surface repellence of the WT biofilms (Fig. 1d). In contrast to the WT strain, the water droplets on the surfaces of *yjbI* disruptant biofilms assumed a contact angle of 30 ± 1.55°, indicating a significant loss of surface repellence in the disruptant biofilms. Similarly, the *tasA* disruptant (COTNd, Supplementary Table 1), which was a negative control lacking TasA, the main fibrous protein in *B. subtilis* biofilms (Diehl et al., 2018, Hobley et al., 2015, Romero et al., 2010), also failed to form biofilms with normal wrinkles and water repellence (Fig. 1d). When the *yjbI* gene was reintroduced into the *yjbI* disruptant, the ability of the strain to form biofilms with wrinkles and surface repellence was markedly recovered (Fig. 1d). These results indicated YjbI is required for normal biofilm formation. The loss of surface repellence in the *yjbI* disruptant biofilm suggests that its biofilm surfaces are wetter than those of WT, which is likely to be related to the lower stiffness phenotype of this mutant strain (Fig. 1a-c).

### YjbI suppresses ROS-mediated TasA aggregation

Since the *yjbI* disruptant lost the biofilm surface repellence in a manner like that of the *tasA* disruptant, we examined the effect of *yjbI* deficiency on TasA. His-tagged TasA (TasA-His) was expressed in the WT and *yjbI* disruptant strains carrying pHtasA1 by cultivating in the biofilm-promoting liquid MSgg medium, and the soluble and insoluble protein fractions from the lysate of the pellicles of each strain were analysed by SDS-PAGE followed by western blotting using an anti–His-tag antibody. A band with the expected molecular mass of 31 kDa for TasA-His was detected in the soluble fractions from both the WT and *yjbI*-disrupted strains (Fig. 2a). The 31-kDa TasA-His band was also detected in the insoluble fraction from the WT strain (Fig. 2a). In contrast, we observed smeared bands with markedly high molecular masses (> 225 kDa) in the insoluble fraction from the *yjbI* disruptant (Fig. 2a), suggesting that YjbI prevents TasA from forming high-molecular-mass aggregates. No monomeric TasA was detected in the insoluble fraction from the *yjbI* disruptant even under the reducing conditions during SDS-PAGE, strongly suggesting the involvement of covalent cross-linking bonds in the formation of the large TasA aggregates in the absence of YjbI (Fig. 2a).

**Fig. 2.**
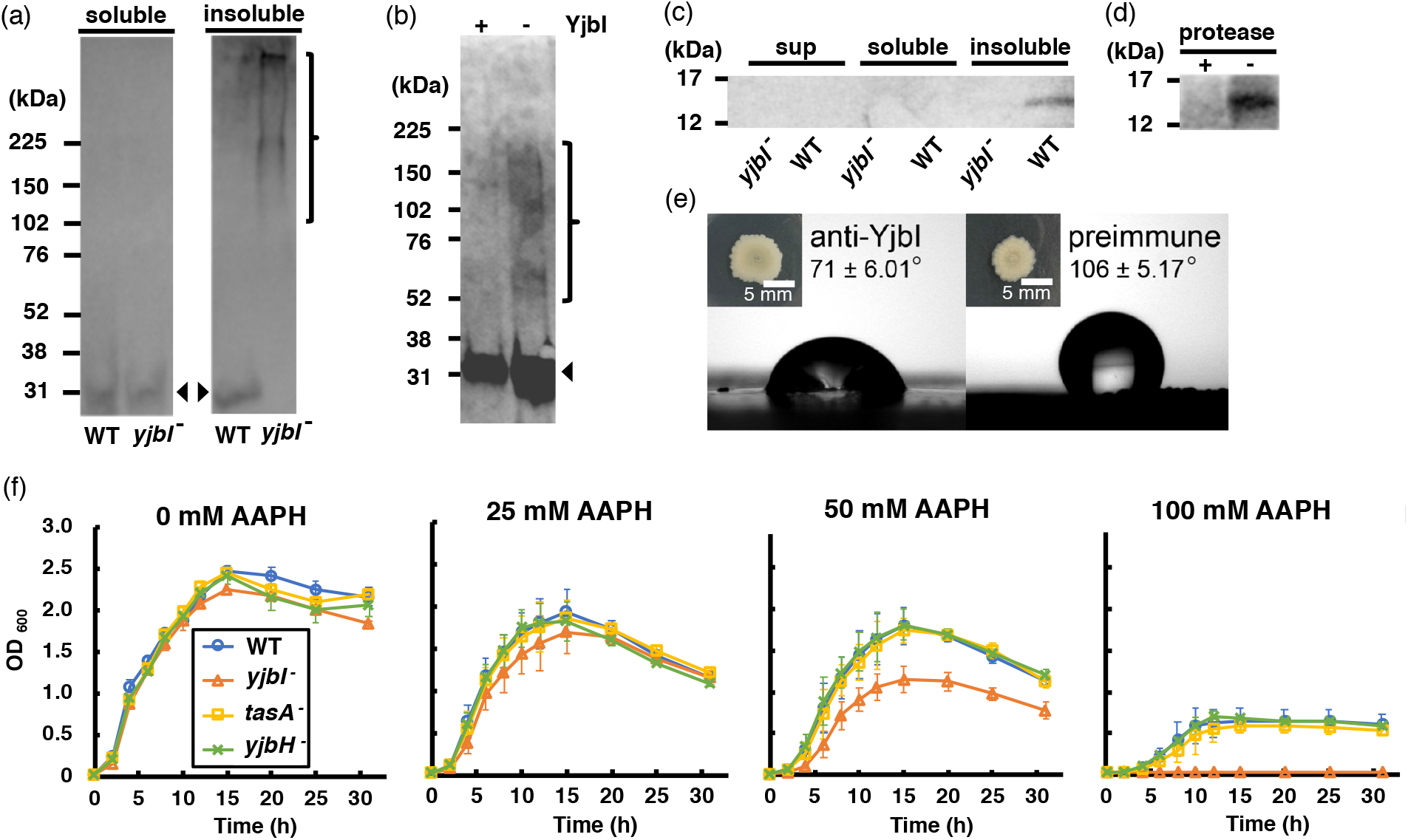
*yjbI* is required to prevent oxidative TasA aggregation and for the normal antioxidant properties of *B. subtilis* cells. (a) Detection of His-tagged TasA (TasA-His) in the soluble (left panel) and insoluble (right panel) fractions of the pellicles of the WT strain or *yjbI* disruptant (*yjbI^-^*) strain carrying pHtasA1, which constitutively expresses TasA-His (Supplementary Table 1). After culturing in liquid MSgg medium, lysates of these pellicles were further analysed by western blotting using an anti-His-tag antibody. The positions of the molecular mass marker proteins are shown on the left. The arrowheads and bracket indicate monomeric and aggregated TasA-His, respectively. Similar results were obtained from three independent experiments. (b) *In vitro* ROS-induced formation of aggregates of purified TasA (pEtasA2-derived TasA_28-261_-His) (Supplementary Table 1) in the presence (+) or absence (-) of purified recombinant YjbI, as evidenced by western blotting analysis using an anti-His-tag antibody. The arrowhead and bracket indicate monomeric and aggregated TasA_28-261_-His, respectively. Protein aggregation was induced with 1 mM H_2_O_2_ and 10 μM FeCl_2_. Similar results were obtained from two independent experiments. (c) Localisation of YjbI in the insoluble biofilm fraction. The soluble and insoluble fractions of the WT and *yjbI^-^* pellicles and culture supernatants (sup) were analysed by western blotting using an anti-YjbI antiserum. Similar results were obtained from two independent experiments. (d) Localisation of YjbI to the cell surface, as evidenced by western blot analysis of the intact WT pellicles with (+) or without (-) protease digestion. YjbI was detected using an anti-YjbI antiserum. Similar results were obtained from two independent experiments. (e) Images showing biofilm surface repellency. Two microliters of rabbit anti-YjbI serum (left panel) or pre-immune serum (right panel) were mixed with 2 μL of pre-cultured *B. subtilis* cells and inoculated on MSgg solid medium. The morphologies of the biofilms are shown in the insets. The water contact angles indicated in the right panels represent the mean ± SD of three independent experiments. (f) Sensitivity of planktonically grown *B. subtilis* strains to AAPH-induced oxidative stress. *B. subtilis* cells (WT, *yjbI^-^, tasA^-^*, or *yjbH^-^*) were inoculated at OD_600_ = 0.02 in LB medium containing 0-100 mM AAPH and grown at 37 °C with shaking. The data are the mean ± SD of three independent experiments. **Fig. 2a source data** **Overall image and raw data of the blotting result.** **Fig. 2b source data** **Overall image and raw data of the blotting result.** **Fig. 2c source data** **Overall image and raw data of the blotting result.** **Fig. 2d source data** **Overall image and raw data of the blotting result.**

Covalent cross-linking of proteins is generally known to be caused by ROS-mediated protein oxidation (Davies, 2016, Gebicki, 1997, Dean et al., 1997). We could partly reproduce the TasA aggregation *in vitro* by oxidising purified TasA_28-261_-His (a mature form of TasA with a C-terminal His-tag) at neutral pH in the presence of hydrogen peroxide and Fe^2+^, which undergo the Fenton reaction (Gebicki, 1997, Welch et al., 2002) to produce ROS (Fig. 2b). We investigated whether YjbI could prevent this oxidation-induced aggregation and found that adding purified YjbI (Fig. S1) to the mixture significantly suppressed the formation of TasA_28-261_-His aggregates (Fig. 2b), consistent with the results of the *in vivo* experiment shown in Figure 1b. These findings suggest YjbI suppresses ROS-mediated TasA aggregation.

### YjbI localises to the surface of *B. subtilis*

TasA localises to the cell surface by the Sec pathway (Romero et al., 2010, Romero et al., 2011). Because YjbI suppressed TasA aggregation *in vivo* (Fig. 2a), it is reasonable to expect that YjbI also localises to the cell surface. To address this hypothesis, the soluble and insoluble fractions of cell lysates from WT and *yjbI*-disrupted *B. subtilis* pellicles were analysed by western blotting using an anti-YjbI antiserum. YjbI was detected only in the insoluble fraction from the WT pellicles (Fig. 2c). We then examined whether YjbI is exposed on the cell surface. Toward this end, we treated intact *B. subtilis* WT pellicles with a protease cocktail before western blot analysis of the whole cells. The protease treatment almost eliminated the immunoreactive YjbI from the intact *B. subtilis* WT pellicles, suggesting that YjbI is localised to the extracellular surface (Fig. 2d). Furthermore, we examined whether YjbI could be recognised by anti-YjbI antibodies, which must be cell impermeable. Treatment of the intact WT cells with the anti-YjbI antiserum before cultivation on MSgg medium impaired colony biofilm formation, with a significant loss of surface repellence (Fig. 2e), as observed for the *yjbI* disruptant. In contrast, treatment with the control pre-immune serum showed almost no effect. These observations suggest that anti-YjbI antibodies can interact with YjbI to impair its function on the biofilm surface. Collectively, these results demonstrate functional YjbI is localised to the extracellular surface to facilitate normal biofilm formation by suppressing ROS-mediated TasA aggregation.

### YjbI functions as an antioxidant protein in *B. subtilis*

To address whether YjbI functions exclusively in biofilm maintenance or in a general cellular protection against oxidative stress, we analysed the effect of 2,2’-azobis(2-amidinopropane) dihydrochloride (AAPH)-induced oxidative stress on the planktonic growth of *B. subtilis* strains in lysogeny broth (LB) liquid medium under shaking conditions. AAPH was chosen as the radical initiator because it gently generates free radicals under neutral pH (Gebicki and Gebicki, 1993) and is suitable for a long-term cultivation experiment. We observed no significant difference between the WT strain and *tasA* disruptant in the sensitivity to AAPH, indicating that TasA plays no apparent role in the protection of the planktonically grown cells from oxidative stress (Fig. 2f). In contrast, the *yjbI* disruptant showed hypersensitivity to AAPH compared with the WT and *tasA* disruptant (Fig. 2f). This result shows that the hypersensitivity of the *yjbI* disruptant to AAPH is not because of TasA impairment in this mutant but presumably because YjbI is involved in a general cellular protection against AAPH-induced oxidative stress, at least during the planktonic growth. Interestingly, disruption of *yjbH* (BKE11550, Supplementary Table 1), which is co-transcribed with *yjbI* (Rogstam et al., 2007), did not alter the sensitivity to AAPH-induced oxidative stress (Fig. 2f), suggesting that YjbI functions independently of YjbH under oxidative stress.

The effect of the oxidative stress induced by hypochlorous acid (HOCl) on the *yjbI* disruptant was also examined. HOCl is a strong bactericidal agent that can cause severe protein hydroperoxidation via the generation of radicals on amino acid residues (Hawkins and Davies, 1998, Stadtman and Levine, 2003). A minimum bactericidal concentration assay revealed that the sensitivity of the *yjbI* disruptant to hypochlorous acid was approximately 100 times higher than the WT strain (Table 1). These data demonstrate that the loss of *yjbI* increases the sensitivity of the cells to oxidative stress, suggesting that YjbI functions as a powerful antioxidant protein in *B. subtilis*. Taken together, these results suggest that the antioxidant role of YjbI is not limited to maintaining the function of TasA in biofilm formation; most likely, YjbI functions to protect any labile cell-surface protein from oxidative damage.

**Table 1.** Minimum bactericidal concentration following exposure to hypochlorous acid (HClO) (n = 2).

|  | HClO (mM) | 62.5 | 31.3 | 15.6 | 5.00 | 2.50 | 1.25 | 0.625 | 0.313 | 0.156 |
| --- | --- | --- | --- | --- | --- | --- | --- | --- | --- | --- |
| 869 | WT | – | + | + | + | + | + | + | + | + |
| 870 | *yjbI*<sup>–</sup> | – | – | – | – | – | – | – | + | + |

(+) visible growth, (–) no visible growth

### YjbI exhibits unprecedented protein hydroperoxide peroxidase-like activity

The observed ROS-mediated TasA aggregation likely involves intermolecular covalent crosslinking among individual TasA proteins (Fig. 2a and 2b). Protein cross-links are formed via radical reactions and the Michael additions after the generation of protein hydroperoxides (halflife: ~4 h at 37 °C) (Davies, 2016), (Gieseg et al., 2000) (Fig. S2). Therefore, we examined *in vitro* whether YjbI could prevent the ROS-mediated aggregation of bovine serum albumin (BSA), which has been a model protein in studies of protein hydroperoxides (Gebicki and Gebicki, 1993). A hydroperoxidised BSA (BSA-OOH) was prepared using the Fenton reaction followed by removal of excess hydrogen peroxide and unstable radicals via cold acetone precipitation (Gieseg et al., 2000). Subsequent incubation of BSA-OOH in the absence of YjbI promoted the spontaneous self-crosslinking of BSA to yield BSA aggregates and a concomitant fragmentation via nonspecific free radical chain reactions (Fig. 3a) (Fig. S2). In contrast, the addition of YjbI to the subsequent incubation of BSA-OOH markedly suppressed the BSA-OOH aggregation (Fig. 3a). An identical result was obtained with BSA-OOH prepared using AAPH (Fig. 3b). Time course analysis further demonstrated the rapid and significant elimination of the hydroperoxide groups from BSA-OOH by YjbI (Fig. 3c). We also tested the effect of adding YjbI before AAPH-induced BSA hydroperoxidisation on the aggregation and fragmentation of BSA. This condition may more closely mimic an *in vivo* environment, since YjbI should coexist with cell-surface proteins, including TasA, before their oxidation. Indeed, pre-addition of YjbI markedly protected BSA from the AAPH-induced aggregation and fragmentation (Fig. 3d). The protective function against oxidative BSA aggregation/fragmentation was specific to YjbI, because two typical haem proteins, haemoglobin, and myoglobin, showed no protective effects, but promoted the oxidative BSA aggregation/fragmentation (Fig. 3d). The effects of haemoglobin and myoglobin are consistent with a previous observation that coexistence of a haem group and an oxidant generally accelerates radical reactions (Svistunenko, 2005). The accumulation of BSA-OOH hydroperoxide groups was also dramatically suppressed by the pre-addition of YjbI (Fig. 3e). Taken together, these results suggest YjbI prevents protein aggregation/fragmentation, most probably through its haem-mediated biologically unprecedented protein hydroperoxide peroxidase-like activity (Fig. S2).

**Fig. 3.**
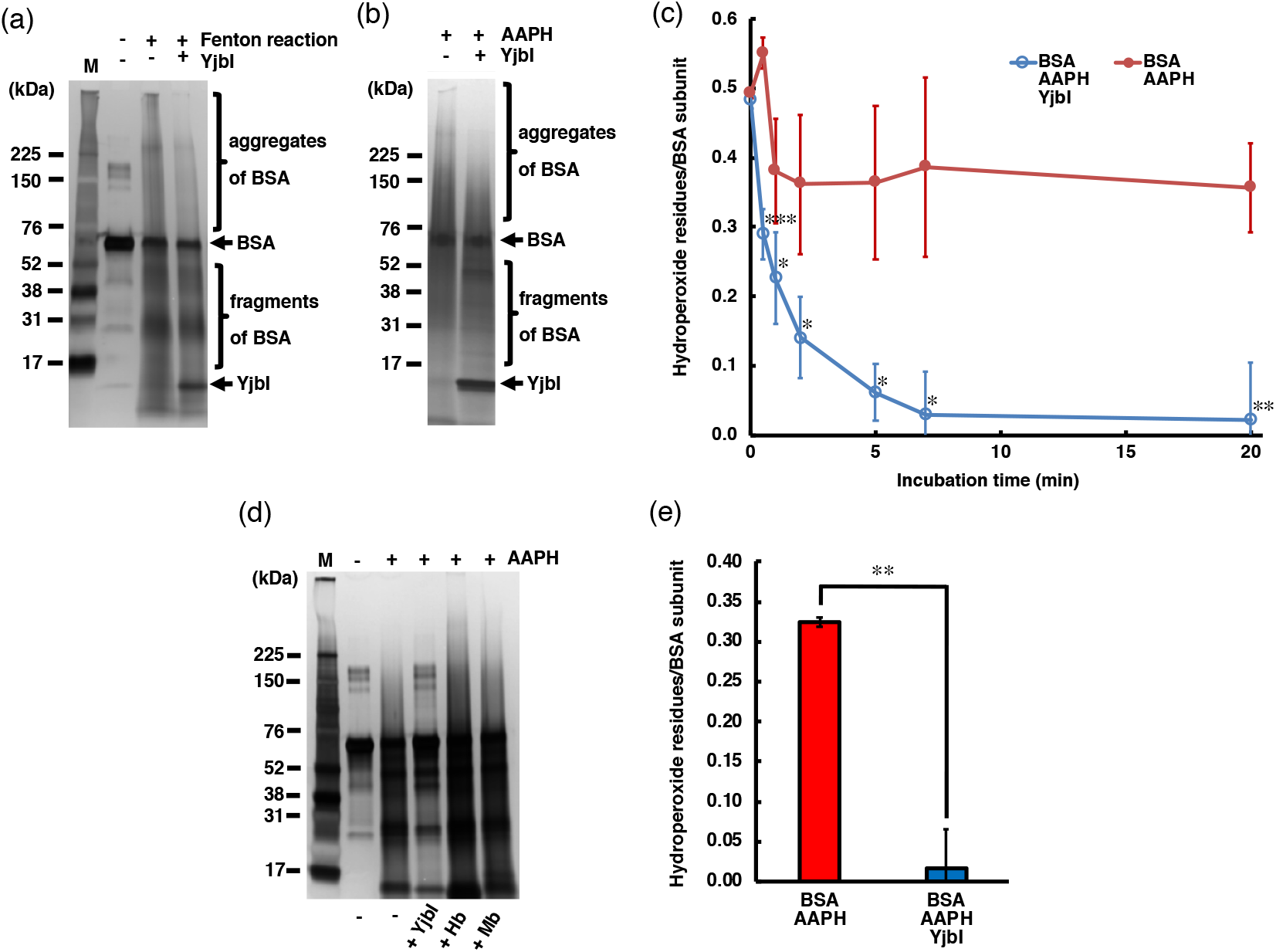
YjbI has a protein hydroperoxide peroxidase activity. The suppressive effect of YjbI on BSA-OOH aggregation induced by the Fenton reaction (a) or AAPH (b). (a) BSA was treated with (+) or without (-) H_2_O_2_ and FeCl_2_ to produce BSA-OOH. BSA-OOH was incubated in the presence (+) or absence (-) of YjbI and analysed by SDS-PAGE and silver staining. YjbI and monomeric BSA are indicated by arrows, and BSA aggregates and fragments are indicated by brackets. The sizes of the molecular mass marker proteins (M) are shown on the left. Similar results were obtained from two independent experiments. (b) BSA-OOH prepared by AAPH treatment was incubated in the presence (+) or absence (-) of YjbI and analysed by SDS-PAGE and silver staining. The labels on the left or right of the images are the same as those in (a). Similar results were obtained from two independent experiments. (c) Time course of the YjbI-induced reduction of the hydroperoxide groups in BSA-OOH. BSA-OOH (30 μM) in 50 mM Tris-acetate buffer (pH 7.0) was incubated with (blue circles) or without (red circles) YjbI (3.3 μM) at 37 °C, and the reaction was terminated by adding 4 volumes of cold acetone at the indicated incubation times (0, 0.5, 1, 2, 5, 7, and 20 min). The number of hydroperoxide groups per BSA subunit was determined for each sample. The data are the mean ± SD of three independent experiments (\**p* < 0.05; \*\**p* < 0.01; and \*\*\**p* < 0.005; *t*-test, one-tailed). (d) YjbI prevents AAPH-induced BSA-OOH aggregation. BSA was incubated for 3 h with (+) or without (-) AAPH before adding YjbI, haemoglobin from bovine blood (Hb) (Sigma-Aldrich), myoglobin from equine heart (Mb) (Sigma-Aldrich), or none (-) and then analysed by SDS-PAGE and silver staining. The positions of the molecular mass marker proteins (M) are shown on the left. Similar results were obtained from two independent experiments. (e) YjbI suppresses AAPH-induced accumulation of BSA-OOH. BSA (6.0 μM) was incubated with 25 mM AAPH in 50 mM Tris-acetate buffer (pH 7.0) with (blue bar) or without (red bar) YjbI (6.6 μM) for 3 h at 37 °C before determining the hydroperoxide groups per BSA subunit. The data are the mean ± SD of three independent experiments (\*\**p* < 0.01; *t*-test, one-tailed). **Fig. 3a source data** **Overall image and raw data of the gel.** **Fig. 3b source data** **Overall image and raw data of the gel.** **Fig. 3d source data** **Overall image and raw data of the gel.**

### YjbI does not affect lipid peroxidation

Lipids, one of the cell-surface components, are also hydroperoxidised by ROS (Cabiscol et al., 2000). We examined whether *yjbI* disruption also affects the amount of lipid hydroperoxides in biofilms. The colony biofilms of the WT and *yjbI* disruptant strains were prepared on solid MSgg medium, and the amounts of lipid hydroperoxide in the collected biofilms were determined. Therefore, the level of lipid hydroperoxide was not significantly increased in the *yjbI* disruptant relative to the level in the WT, implying that YjbI does not function in eliminating lipid hydroperoxides (Fig. S3).

## Discussion

In this study, we revealed a physiological and biochemical function of YjbI. Our results suggest YjbI prevents the ROS-induced aggregation of proteins, such as TasA, localised on the cell surface of *B. subtilis*. We also showed that YjbI may possess a unique protein hydroperoxide peroxidase-like activity, absent from other haem proteins, such as haemoglobin and myoglobin. Oxidative protein aggregation and protein carbonyls derived from protein radicals and hydroperoxides (Davies, 2016) have emerged as important biomarkers of various cellular defects caused by oxidative stress not only in mammals (Korovila et al., 2017, Heinecke et al., 1993) but also in bacteria (Ling et al., 2012).

In *B. subtilis*, several cytosolic factors have been known to protect the cells from oxidative damages. The catalase KatA (Chen et al., 1995) and hydrogen peroxide peroxidase AhpC (Broden et al., 2016) protect the cells from hydrogen-peroxide–induced oxidative damages. OhrA and OhrB contribute to the cellular protection from organic hydroperoxides (Fuangthong et al., 2001). In addition, bacillithiol has been proposed to be involved in superoxide stress and metal homeostasis, but not in hydrogen-peroxide-induced oxidative stress (Fang and Dos Santos, 2015, Gaballa et al., 2010). However, much less is known about the mechanism protecting cell-surface proteins from oxidative damages in gram-positive bacteria. In the gram-positive *Streptococcus pneumoniae*, surface-exposed proteins with methionine sulfoxide residues are reduced by the membrane-bound methionine sulfoxide reductase MsrAB2 (Saleh et al., 2013), but the gene of an equivalent membrane-anchored enzyme is absent from the *B. subtilis* genome. To our best knowledge, YjbI is the first example of an antioxidant protein involved in protecting cell-surface proteins against oxidative aggregation. Antioxidation of cell-surface proteins is important, especially for proteins located and exposed at the air-liquid interface, such as the biofilm surface (Beloin et al., 2004, Ezraty et al., 2017). Disrupting *yjbI* had a crucial effect on the integrity of the biofilm cell surface, presumably because of this reason (Fig. 1).

*M. tuberculosis* trHbO has an autokinase activity and plays a role in the survival and adaptation of the bacterium under hypoxia (Hade et al., 2020). However,*M. tuberculosis*trHbO shares a moderate (31%) amino acid sequence identity with *B. subtilis* YjbI, and *M. tuberculosis* carries another truncated haemoglobin, trHbN (Couture et al., 1999), which is missing in *B. subtilis*. Moreover, because *M. tuberculosis* is an obligate aerobic bacterium, its responses under oxygen stress may be largely different from those of the member of the facultative anaerobes *B. subtilis*. Whether YjbI orthologues found in other bacteria function in the suppression of oxidative protein aggregation remains an open question for future research.

In this study, we demonstrated YjbI reduces hydroperoxide groups in BSA-OOH via its peroxidase-like activity and suppressed the spontaneous aggregation of BSA-OOH. Typical haem peroxidases, such as horseradish peroxidase and lactoperoxidase, exhibit negligible activity on protein hydroperoxides (Davies, 2016, Morgan et al., 2004, Gebicki et al., 2002). Haemoglobin and myoglobin can react with small peptide-sized hydroperoxides but not with large protein hydroperoxides, such as BSA-OOH (Morgan et al., 2004). The X-ray crystal structure of *B. subtilis* YjbI (PDB ID: 1UX8) (Giangiacomo et al., 2005) shows this protein has a 55-Å^2^ surface opening (Fig. S4), which may allow direct access of bulk solvent and large molecules to the haem active-site. Therefore, the structural feature probably confers the potential to react with a protein hydroperoxide on YjbI.

An electron donor is required for the proposed protein hydroperoxide peroxidase-like reaction by YjbI. We showed that the *in vitro* reduction of BSA-OOH by YjbI proceeded without addition of any external electron donors (Fig. 3). This observation implies that electrons needed for the *in vitro* protein hydroperoxide peroxidase-like reaction may be provided from the amino acid residues of YjbI itself or BSA. Interestingly, in the hydrogen peroxide peroxidase reaction catalysed by *M. tuberculosis* trHbO, the tyrosine residues near the haem of trHbO have been suggested to serve as electron donors (Ouellet et al., 2007). At least two of these tyrosine residues are conserved in trHbO orthologues (Tyr25 and Tyr63 of YjbI), which are located close to the haem according to the results of the crystal structure analysis (Giangiacomo et al., 2005). Indeed, Tyr can be stabilised by providing electrons to yield either dityrosine or a series of oxidised derivatives of dityrosine, 3,4-dihydroxyphenylalanine (DOPA), DOPA semiquinone, and DOPA quinone (Maskos et al., 1992, Giulivi and Davies, 2001). Taken together, we propose that the Tyr residues in YjbI are the most promising candidates for donating electrons for the *in vitro* YjbI-catalysed protein hydroperoxide peroxidase-like reaction. Nevertheless, it remains unclear whether YjbI utilises an endogenous or exogenous electron donor other than the amino acid residues of YjbI itself *in vivo*. Previous reports have shown that *M. tuberculosis* trHbO can receive electrons from ferrocytochrome *c* (Ouellet et al., 2007). Alternatively, YjbH is a DsbA-like protein (Guddat et al., 1998) with two active-site cysteine residues that may have a potential to donate electrons for the YjbI-catalysed reduction of protein hydroperoxides (Fabianek et al., 2000). However, the role of YjbH as an electron donor to YjbI is unlikely because disrupting *yjbH* did not affect the sensitivity to AAPH (Fig. 2f), and YjbH localises to the cytosol, unlike YjbI, to function as an effector of Spx, a central regulator of stress response (Larsson et al., 2007).

The localisation of YjbI to the cell surface is consistent with its role in protecting TasA and other cell-surface proteins from ROS-induced aggregation (Fig. 4a and 4b). However, there is no apparent targeting signal sequence in the amino acid sequence of YjbI, and bioinformatic analysis using SignalP (http://www.cbs.dtu.dk/services/SignalP/abstract.php) or PSORT (https://www.psort.org) predicted that YjbI is a cytoplasmic protein (Fig. S5). Purified recombinant YjbI, which was produced in *E. coli,* was obtained in a soluble form (Fig. S1), in contrast to the insoluble feature of the native YjbI in *B. subtilis*. Analogously, *M. tuberculosis* trHbN belonging to another group of trHb family, is also devoid of a targeting signal sequence, is localised to the cell surface following glycosylation, whereas the corresponding recombinant protein expressed in *E. coli* is detected in a soluble form (Arya et al., 2013). Nevertheless, no glycosylation site is found on YjbI. Therefore, the mechanism for the translocation of YjbI to the extracellular surface should be investigated in future studies.

**Fig. 4.**
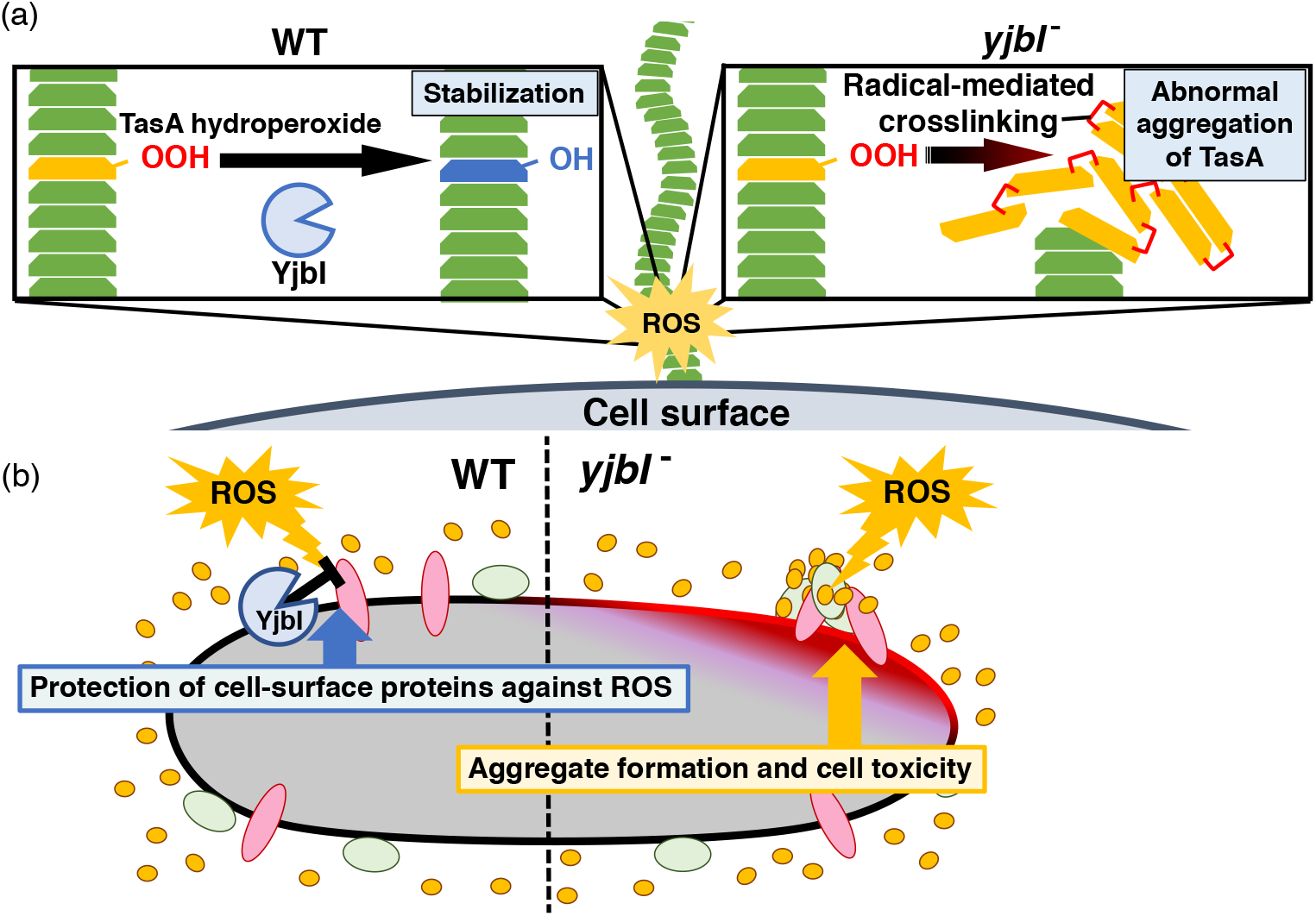
YjbI protects proteins and the cell from reactive oxygen species (ROS) by removing hydroperoxide groups from proteins. (a) YjbI repairs the oxidatively damaged TasA-OOH in biofilms. In the wild-type (WT) strain (left panel), the normal fibrous TasA (green trapezoids) is damaged by ROS to generate TasA-OOH (yellow trapezoid) on the cell surface. YjbI converts TasA-OOH to TasA-OH (blue trapezoid) to stabilise the normal TasA fibre. In contrast, TasA-OOH aggregates via radical-mediated protein cross-linking in the *yjbI^-^* strain (right panel). (b) A proposed role of YjbI in the general protection of cell-surface proteins from ROS-induced oxidative damage. Protein hydroperoxide-modified residues generated in various cellular surface proteins (red, green, and yellow ellipses) are generally reduced to protein hydroxy residues by YjbI to protect from further damage (i.e., aggregation) (left half). *yjbI* disruption accumulates protein aggregates via hydroperoxidation and cross-linking of various proteins, resulting in cell toxicity (right half).

Although protein hydroperoxides are one of the major products formed because of ROS-mediated protein oxidation (Davies et al., 1995, Gebicki, 1997), (Gebicki and Gebicki, 1999), little is known about the effects of their generation on bacterial physiology. Biofilm matrix proteins are expected to be exposed to an air-liquid interface, which is a region with a relatively high risk of protein oxidation. Therefore, repairing the oxidatively damaged proteins on the cell surface may be crucial for bacterial survival in biofilms. The loss of biofilm integrity associated with the irreversible aggregation of the biofilm matrix protein TasA in the *yjbI*-disrupted strain may provide an important clue to further understand the significance of protein hydroperoxide generation on the cell surface.

The findings of this study may lead to future applications. An increasing number of studies have suggested that biofilms are associated with bacterial infections in humans and with natural resistance of pathogenic bacteria to antibiotics (Hall and Mah, 2017). We found that *B. subtilis* biofilm formation can be suppressed by an antiserum against YjbI (Fig. 1f) and that YjbI deficiency severely impairs the bacterial tolerance to the oxidative stress induced by AAPH (Fig. 2d) or HOCl (Table 1). HOCl oxidises various amino acid residues such as methionine and lysine. In addition, the reaction of HOCl with O_2_ helps to form hydroxy radicals (Candeias et al., 1993, Candeias et al., 1994), which initiates protein peroxidation (Stadtman and Levine, 2003). Intriguingly, leukocytes employ ROS, including peroxides and HOCl, in the respiratory burst during the innate immune response to bacterial infection (Babior, 1984). Therefore, the YjbI orthologues in pathogenic bacteria would be potential novel drug targets to inhibit pathogenic bacterial growth and biofilm formation. In fact, trHbO is implicated in *Mycobacterium* pathogenicity (Wittenberg et al., 2002). Future studies on the structurefunction relationship of YjbI may contribute to developing a specific inhibitor that suppresses infections by the wide variety of trHbO-harbouring pathogenic bacteria (e.g., *Bacillus, Mycobacterium*, and *Staphylococcus*).

## Methods

### Bacterial strains, plasmids, and culture conditions

The bacterial strains and plasmids used in this study are listed in Supplementary Table 1. *B. subtilis* 168 and *E. coli* derivatives were obtained by transformation of competent cells with plasmids according to standard protocols (Harwood and Cutting, 1990, Studier and Moffatt, 1986). For pre-cultivation, *B. subtilis* strains were grown in LB medium at 37 °C for 8 h. Recombinant *E. coli* strains were cultivated in LB medium at 37 °C. When appropriate, 30 μg/mL tetracycline and 100 μg/mL kanamycin were added to the cultures of *B. subtilis* and *E. coli*, respectively.

### Pellicles and colony biofilm formation

Pellicles in liquid media were prepared by inoculating 1/20 volume of the *B. subtilis* pre-culture in the biofilm-promoting MSgg medium (Bucher et al., 2016) (5 mM potassium phosphate in 100 mM MOPS buffer at pH 7.0 supplemented with 2 mM MgCl_2_, 700 μM CaCl_2_, 50 μM MnCl_2_, 50 μM FeCl_3_, 1 μM ZnCl_2_, 2 μM thiamine, 0.5% (w/w) glycerol, 0.5% (w/w) glutamate, 0.005% (w/w) phenylalanine, 0.005% (w/w) tryptophan, and 0.005% (w/w) threonine), followed by cultivation at 37 °C for 48 h without shaking. Bacterial colony biofilms on a solid medium were obtained by inoculating a 3-μL aliquot of the *B. subtilis* pre-culture on solid MSgg medium containing 1.5% (w/w) agar, followed by cultivation at 37 °C for 48 h.

### AFM analysis of biofilms

A *B. subtilis* pellicle was gently placed on a silicon substrate for surface analysis by AFM. All measurements were carried out using a scanning probe microscope (E-Sweep; Hitachi High-Tech Science Corporation, Tokyo, Japan) equipped with a microcantilever (SI-DF3, Hitachi High-Tech Science Corporation) with a spring constant of 1.5 N/m and a resonance frequency of 26 kHz. The relative surface softness and the length of the long axis of 50 individual bacterial cells in an AFM image were analysed for each strain using ImageJ (NIH, v.1.52k) (https://imagej.nih.gov/ij/).

### Biofilm surface repellence analysis

Biofilm surface repellence was evaluated according to a described method (Arnaouteli et al., 2017) by measuring the contact angle of a 2-μL water droplet placed on the centre of a bacterial colony biofilm formed on solid MSgg medium, using a DSA100 drop shape analyser (KRÜSS).

The water droplet was equilibrated at 28 °C for 5 min before imaging and measurement. The contact angle is presented as the mean ± SEM of at least three independent experiments.

### Fractionation of soluble and insoluble *B. subtilis* biofilm proteins

*B. subtilis* pellicle (0.1 g) was harvested using a spatula and lysed by incubation with 0.5 mL of B-PER saturated with phenylmethylsulfonyl fluoride (PMSF, Sigma-Aldrich), supplemented with 5 mg/mL lysozyme at 28 °C for 2 h, and centrifuged. Both the soluble proteins in the supernatant and the precipitated insoluble proteins were recovered and used for further studies.

### Purification of YjbI and TasA produced in *E. coli*

To purify recombinant YjbI, *E. coli* BL21(DE3)/pEyjbI2 cells (Supplementary Table 1) were grown in LB medium at 37 °C for 6 h, followed by induction of gene expression with 25 μM isopropyl β-D-1-thiogalactopyranoside and further cultivation for 10 h. After centrifugation, harvested cells were lysed in B-PER (Thermo Fisher Scientific) supplemented with approximately 1 mM PMSF according to the manufacturer’s instructions (Thermo Fisher Scientific). After centrifugation, the crude extract was fractionated with ammonium sulphate (30–60% saturation) for 16 h at 4 °C. After centrifugation, the protein precipitate was resuspended in 50 mM MOPS buffer (pH 7.0) and then applied to a Sephacryl S-100 column (GE Healthcare) equilibrated with the same buffer. Red haemoprotein fractions from the column were collected and passed through an Amicon Ultra 100 kDa device (Millipore). The filtrate containing YjbI was applied to a Toyopearl DEAE-650M column (Tosoh) equilibrated with 50 mM Tris-acetate buffer (pH 7.4). YjbI was eluted at a flow rate of 0.25 mL/min with a linear NaCl gradient (0–0.5 M) prepared in the same buffer. The fractions containing YjbI were desalted and concentrated using an Amicon Ultra 10 kDa device in 50 mM Tris-acetate buffer (pH 7.4) to yield the purified YjbI preparation.

TasA_28-261_-His, a mature form of TasA (Diehl et al., 2018) (amino acid residues 28-261) with a His-tag at the C-terminus was produced in *E. coli* BL21(DE3)/pEtasA2 (Supplementary Table 1) by culturing the cells in LB medium at 37 °C for 6 h, followed by induction with 25 μM isopropyl β-d-1-thiogalactopyranoside and further cultivation for 10 h. The cells were harvested by centrifugation and lysed in B-PER supplemented with about 1 mM PMSF according to the manufacturer’s instructions. The crude extract was applied onto a nickel chelation column (His GraviTrap, GE Healthcare), and TasA_28-261_-His was eluted according to the manufacturer’s instructions.

### *In vitro* ROS-induced TasA aggregation

Purified TasA_28-261_-His (200 μg) was incubated with 1 mM hydrogen peroxide and 10 μM FeCl_2_ in the presence or absence of YjbI (15 μg) in 50 mM Tris-acetate buffer (pH 7.4) at 37 °C for 30 min. The proteins, after being precipitated by adding four volumes of cold acetone, were harvested by centrifugation, and dissolved in a standard SDS sample buffer (0.125 M Tris (pH 6.8), 10% (w/w) 2-mercaptethanol, 4% (w/w) SDS, 10% (w/w) sucrose, 0.01% (w/w) bromophenol blue), which was followed by SDS-PAGE and western blot analysis of TasA_28-261_-His with an anti–His-tag antibody.

### Analysis of YjbI cell-surface localisation following extracellular protease digestion

YjbI cell-surface localisation was analysed following extracellular protease digestion of an intact *B. subtilis* pellicle. *B. subtilis* pellicle (10 mg) was incubated in 50 mM Tris-acetate buffer (pH 7.4) containing 5 mg/mL protease mixture (Pronase E, Sigma-Aldrich) in a total volume of 1 mL at 28 °C for 2 h and then directly resuspended in the standard SDS sample buffer before SDS-PAGE and western blot analysis.

### SDS-PAGE and western blot analysis

Lysates in the standard SDS sample buffer were heated and then resolved on a 4–20% SDS/PAGE gradient gel (Mini-PROTEAN, Bio-Rad), followed by electroblotting on a PVDF membrane via a Trans Blot Turbo (Bio-Rad) at 25 V for 30 min. The membrane was blocked with 1% (w/w) skim milk in TBS (50 mM Tris-HCl and 0.85% (w/w) NaCl, pH 7.4) at 37 °C for 1 h, incubated with an anti-YjbI or anti-His-tag antibody at a dilution of 1: 1000 (vol/vol) in 37 °C TBS for 2 h, washed in TBST (TBS with 0.1% (v/v) Tween-20), and then incubated with a secondary goat anti-rabbit (for the anti-YjbI antiserum) or goat anti–mouse (for the anti–His-tag antibody) antibody conjugated with horseradish peroxidase (Bio-Rad) for 2 h. After the membrane was washed in TBST, the immunoreactive bands were visualised using 1-Step Ultra TMB (Pierce). The membrane was photographed using a gel imager (Gel Doc, Bio-Rad). The anti-His-tag monoclonal antibody was purchased from Medical and Biological Laboratories (Nagoya, Japan), and the anti-YjbI antiserum was a rabbit serum purchased from Sigma Genosys. The anti-YjbI antiserum was prepared by immunising rabbits with purified YjbI mixed with complete Freund’s adjuvant, boosting five times with each a week apart.

### Planktonically grown *B. subtilis* strains under AAPH-induced oxidative stress

After growth in LB medium at 37 °C for 8 h, 3-μL LB medium suspensions of *B. subtilis* strains were aliquoted and cultured on solid LB medium with 0, 25, 50, or 100 mM AAPH at 37 °C for 31 h. A small amount of each culture was fractionated at 0, 2, 4, 6, 8, 10, 12, 15, 20, 25, and 31 h. Next, each fractionated culture was diluted with 0.9% (w/w) saline to keep the absorbance ≤ 1.0 and then measured at 600 nm.

### Minimum bactericidal concentration assay for hypochlorous acid

Pre-cultured *B. subtilis* strains (50 μL) were mixed with 950 μL of phosphate-buffered saline (137 mM NaCl, 8.1 mM Na_2_HPO_4_, 2.68 mM KCl, and 1.47 mM KH_2_PO_4_, pH 7.4) containing HClO at final concentrations between 0 and 500 mM. After incubation at 37 °C for 1 h, a 10-μL aliquot from each mixture was spread on LB solid medium. The cells were grown at 37 °C for 48 h before bacterial colony counting. Then, the minimum concentration of HClO that generated no colony was determined.

### Generation of BSA protein hydroperoxides

BSA-OOH was generated by the Fenton reaction or oxidation with AAPH (Gieseg et al., 2000). In the Fenton reaction, 0.2 mg/mL BSA (fatty-acid-free, Sigma-Aldrich) was incubated with 150 mM hydrogen peroxide and 1 mM FeCl_2_ in 150 mM acetate buffer (pH 5.0) in a total volume of 30 μL at 37 °C for 15 min. To remove excess hydrogen peroxide and iron, 1 mL of 80% cold acetone was added, and the sample was centrifuged at 20,000 × *g* for 10 min to precipitate BSA-OOH. For oxidation with AAPH, 0.2 mg/mL BSA was incubated with 25 mM AAPH in 50 mM Tris-acetate buffer (pH 7.0) in a total volume of 30 μL at 37 °C for 12 h. The reaction was terminated by adding 4 volumes of cold acetone. After centrifugation, the precipitated BSA-OOH was recovered.

### Analysis of the YjbI-mediated prevention of ROS-induced protein aggregation

The effect of YjbI on ROS-induced BSA aggregation was examined via two experiments—a two-step reaction and a one-pot reaction. In the two-step reaction, BSA-OOH was prepared beforehand as described above and then incubated with 0.1 mg/mL YjbI in 50 mM Tris-acetate buffer (pH 7.0) at 37 °C for 3 h in a total volume of 15 μL. In the one-pot reaction, each 0.1 mg/mL globins (YjbI, Hb, or Mb) and 25 mM AAPH was simultaneously added to 0.2 mg/mL BSA in 50 mM Tris-acetate buffer (pH 7.0) in a total volume of 30 μL and incubated at 37 °C for 3 h. For both experiments, the reaction was stopped by adding 4 volumes of ice-cold acetone, and the precipitated proteins were recovered by centrifugation. The proteins were resolved by SDS-PAGE and visualised by silver staining (Wako, Silver Stain II Kit).

### Determination of the hydroperoxide groups in BSA

After hydroperoxidation, the acetone-precipitated BSA was dissolved in 25 mM H_2_SO_4_ to assess for the hydroperoxide groups on the protein, as described (Gieseg et al., 2000). In brief, nine volumes of dissolved BSA-OOH were mixed with one volume of 5 mM ferrous ammonium sulphate and 5 mM xylenol orange in 25 mM H_2_SO_4_ and incubated at 28 °C for 30 min in the dark. The *A*_560_ of the mixture was measured using a UV-Vis spectrophotometer (SH-9000, Corona Electric). The molar extinction coefficient of the xylenol orange-Fe^3+^ complex (1.5 × 10^4^ M^-1^cm^-1^) (Gieseg et al., 2000) in 25 mM H_2_SO_4_ at 28 °C was used to determine the hydroperoxide concentration. The number of hydroperoxide residues per BSA subunit was determined by calculating the mole of peroxide generated per mole of added BSA (molecular weight: 66,463).

#### Statistical analyses

Statistical differences between two experimental groups were identified using one-tailed Student’s *t*-test assuming equal variance (Microsoft Excel for Mac v. 16.16.21). No data points were removed from the data set before the analyses.

## Abbreviations

ROS: reactive oxygen species;
trHb: truncated haemoglobin;
GSH: glutathione;
EPS: exopolysaccharide;
NO: nitric oxide;
c-PTIO: 2-(4-carboxyphenyl)-4,4,5,5-tetramethylimidazoline-1-oxyl-3-oxide;
BSA: bovine serum albumin;
BSA-OOH: hydroperoxidised BSA;
AAPH: 2,2’-azobis(2-amidinopropane) dihydrochloride;
AFM: atomic force microscopy.

## Acknowledgements

The authors thank K. Kitayama and T. Kurihara for help with amino acid sequence analysis. The gene-disrupted strains used in this research were provided by the National BioResource Project (NBRP), Japan. This work was supported by JSPS KAKENHI grant numbers JP18K14383 and JP20K15446, by the Program for the Third-Phase R-GIRO Research from Ritsumeikan University, and by a grant from the Japan Foundation for Applied Enzymology.

## Author contributions

T.I. conceived the idea and performed most experiments. T.I., R.T., J.K., and H.M. designed the experiments. M.T. and K.H. analysed the data from the culture with AAPH and AFM, respectively. T.I. and H.M. wrote the paper.

## Supplementary information

**Supplementary Fig. S1.**
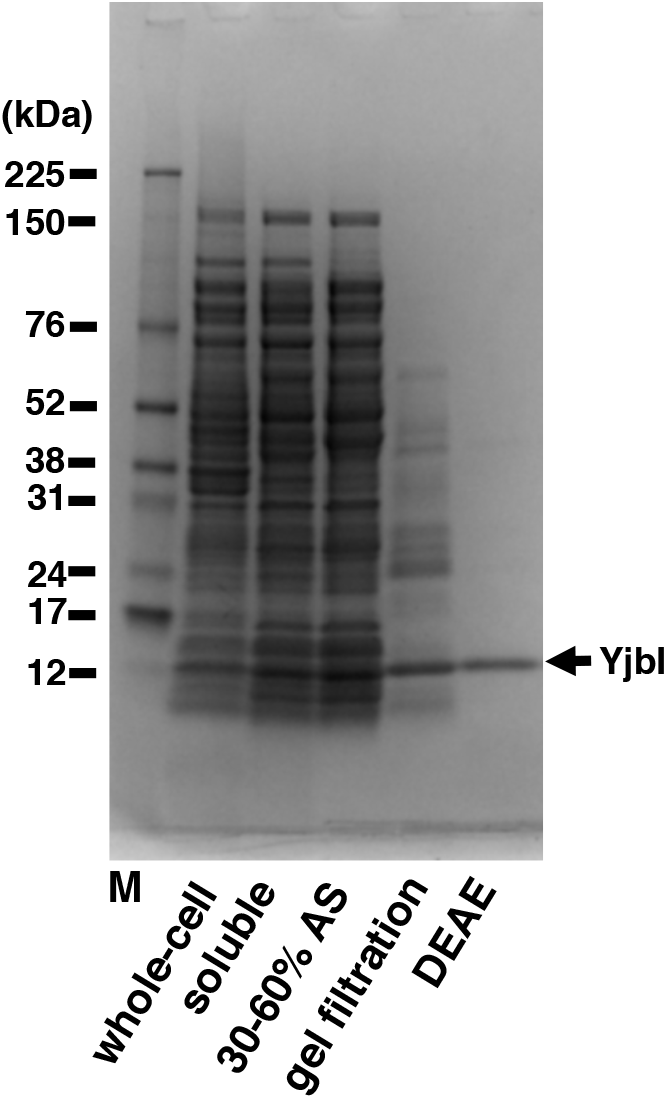
Purification of recombinant YjbI. YjbI was produced in *E. coli* B L21(DE3)/pEyjbI2 cells grown in lysogeny broth (LB) medium. Proteins from each purification step (i.e., whole-cell and soluble crude extracts); 30-60% saturation ammonium sulphate (AS) fractionation; Sephacryl S-100 gel filtration; and Toyopearl DEAE-650M chromatography were resolved by SDS-PAGE and then stained with Coomassie Brilliant Blue. The positions of the molecular mass marker proteins (M) are shown on the left. **Fig. S1 source data** **Overall image and raw data of the gel.**

**Supplementary Fig. S2.**
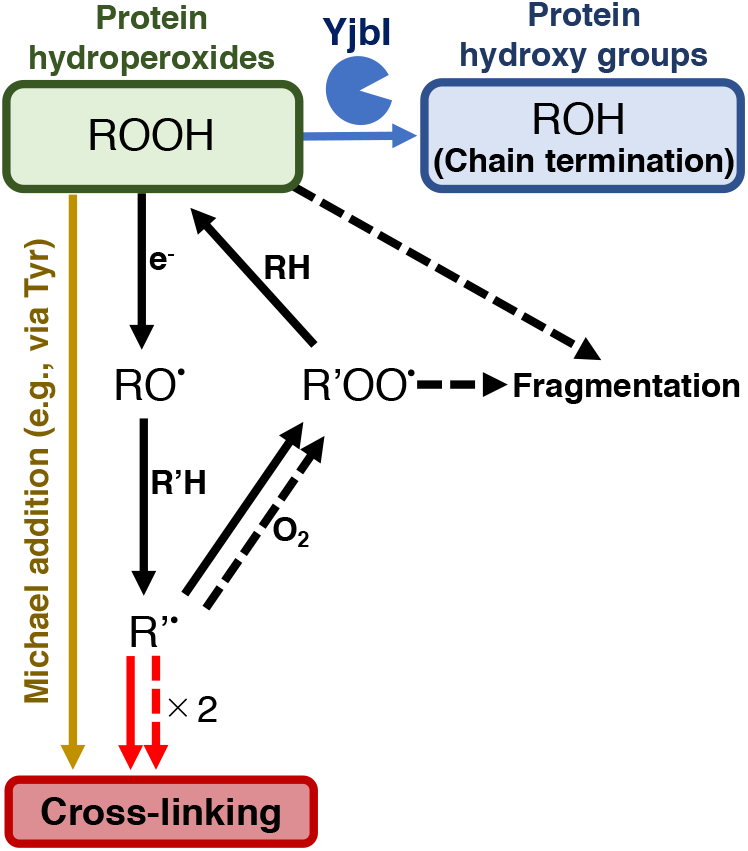
Schematic drawing summarising the proposed protein cross-linking and fragmentation induced by protein radicals derived from protein hydroperoxides and prevention of these reactions by the protein hydroperoxide peroxidase activity of YjbI. The solid and dashed arrows indicate the already known reactions (Davies, 2016) occurring at protein side chains and backbones, respectively. Protein cross-linking proceeds via spontaneous radical coupling (red arrows), and the Michael addition occurs at protein side chains (yellow arrow). YjbI catalyses the protein hydroperoxide peroxidase reaction (blue arrow) to convert protein hydroperoxides (ROOH) to hydroxy groups (ROH) at protein side chains and prevents further protein cross-linking and fragmentation. Side chain alkoxyl radicals (RO·) can be formed through one-electron reduction (e.g., homolysis) of ROOH, followed by hydrogen atom abstraction reactions with other protein side chains or backbones (R’H). The carbon-centred radicals (R’·) of side chains or backbones lead to protein cross-linking to yield protein aggregates (indicated with solid and dashed red arrows). The side chain or backbone R’· also reacts with oxygen to produce protein peroxy radicals (R’OO·) under aerobic conditions. The side chain R’OO· reacts with the side chains or backbones of other amino acid residues (RH) in its vicinity to yield protein ROOH moieties. The unstable backbone R’OO· and ROOH moieties cause protein fragmentation.

**Supplementary Fig. S3.**
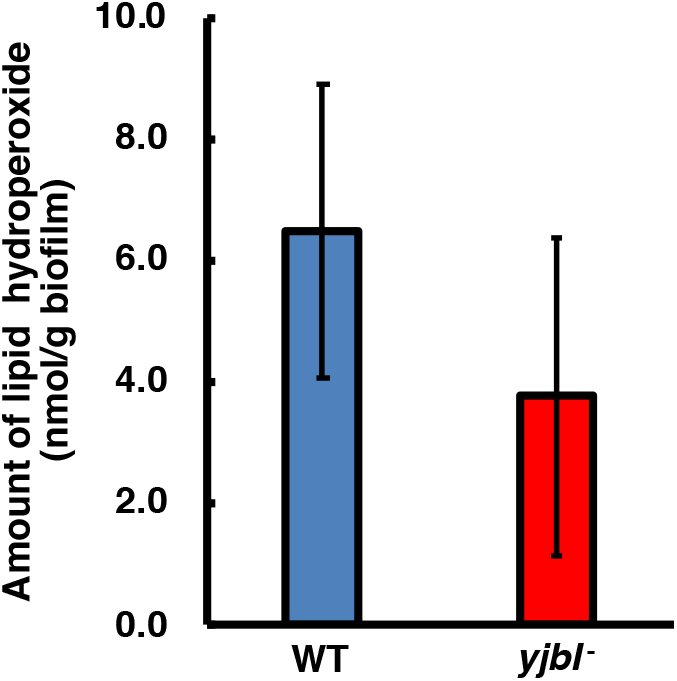
Lipid hydroperoxide (LOOH) levels were not affected in *yjbI*^-^ strain biofilms. Bacterial colony biofilms were obtained by inoculating a 3-μL aliquot of pre-cultured *B. subtilis* on solid MSgg medium, followed by cultivation at 37 °C for 48 h. The biofilms were collected and suspended in phosphate-buffered saline by using a spatula. These WT and *yjbI* disruptant biofilm suspensions were analysed with a colorimetric LOOH assay kit (LPO assay kit, Cayman Chemical Company) according to the manufacturer’s instructions. The data are shown as the mean ± SD of three independent experiments (*p* = 0.083; *t*-test, one-tailed).

**Supplementary Fig. S4.**
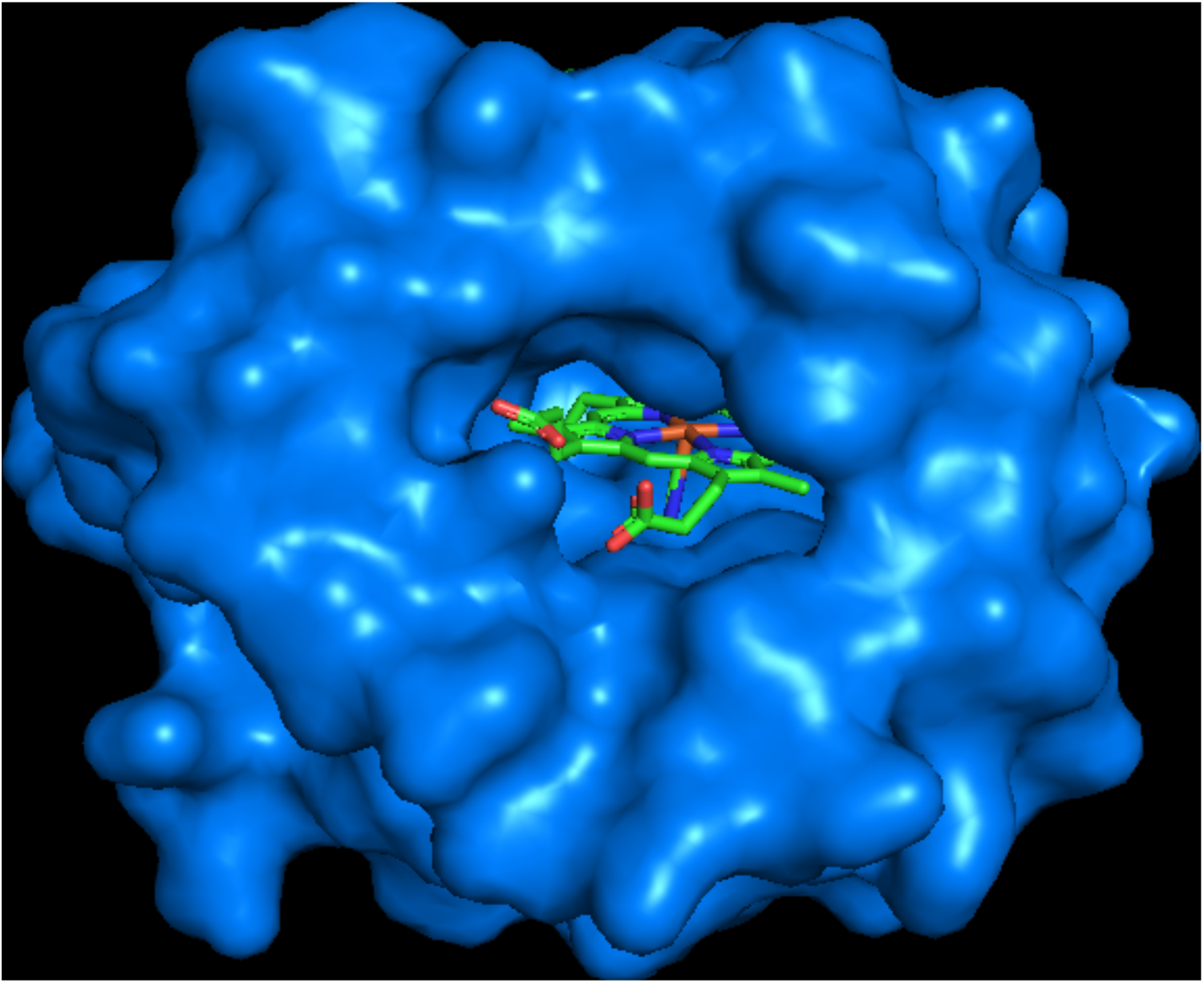
X-ray crystal structure of *B. subtilis* YjbI (PDB ID: 1UX8) (Giangiacomo et al., 2005), showing an opening on the molecular surface. The protein surface is shown in blue, and the haem molecule is shown with a stick model. The image was drawn using PyMOL.

**Supplementary Fig. S5.**
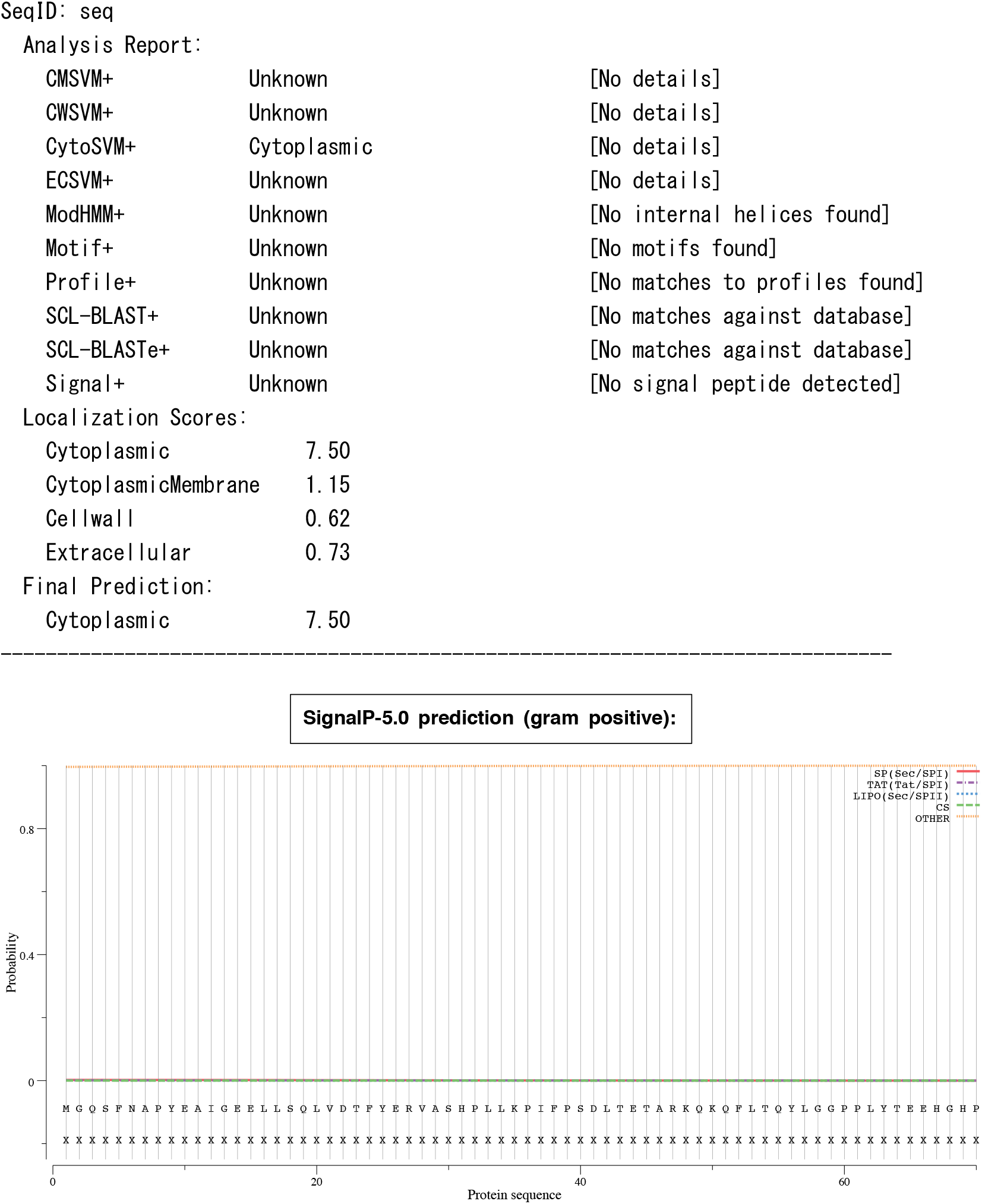
*B. subtilis* YjbI does not contain a predicted signal sequence. The results of bioinformatic analyses by using PSORTb (https://www.psort.org/psortb/) or SignalP-5.0 (http://www.cbs.dtu.dk/services/SignalP/).

**Supplementary Table 1.**
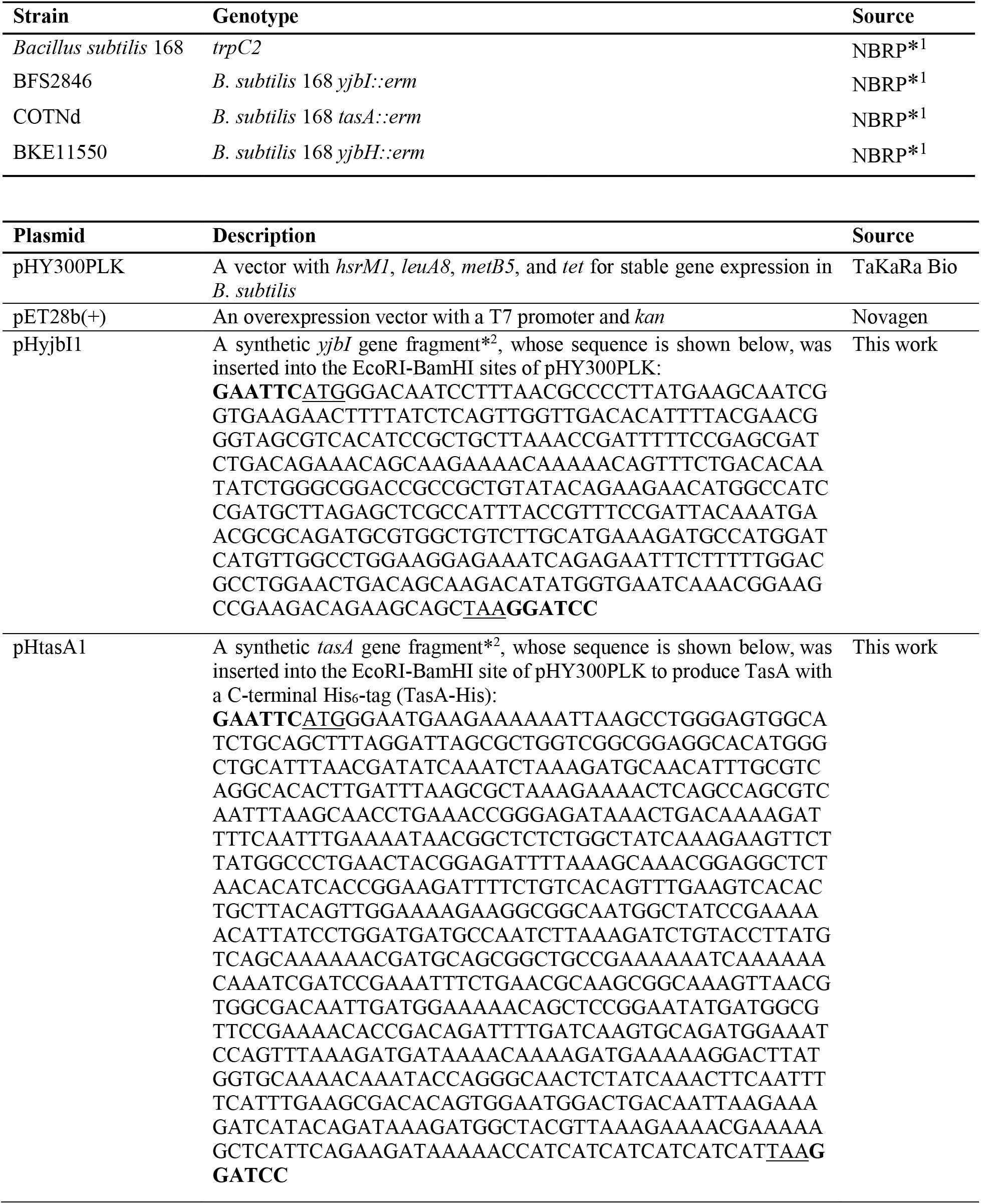
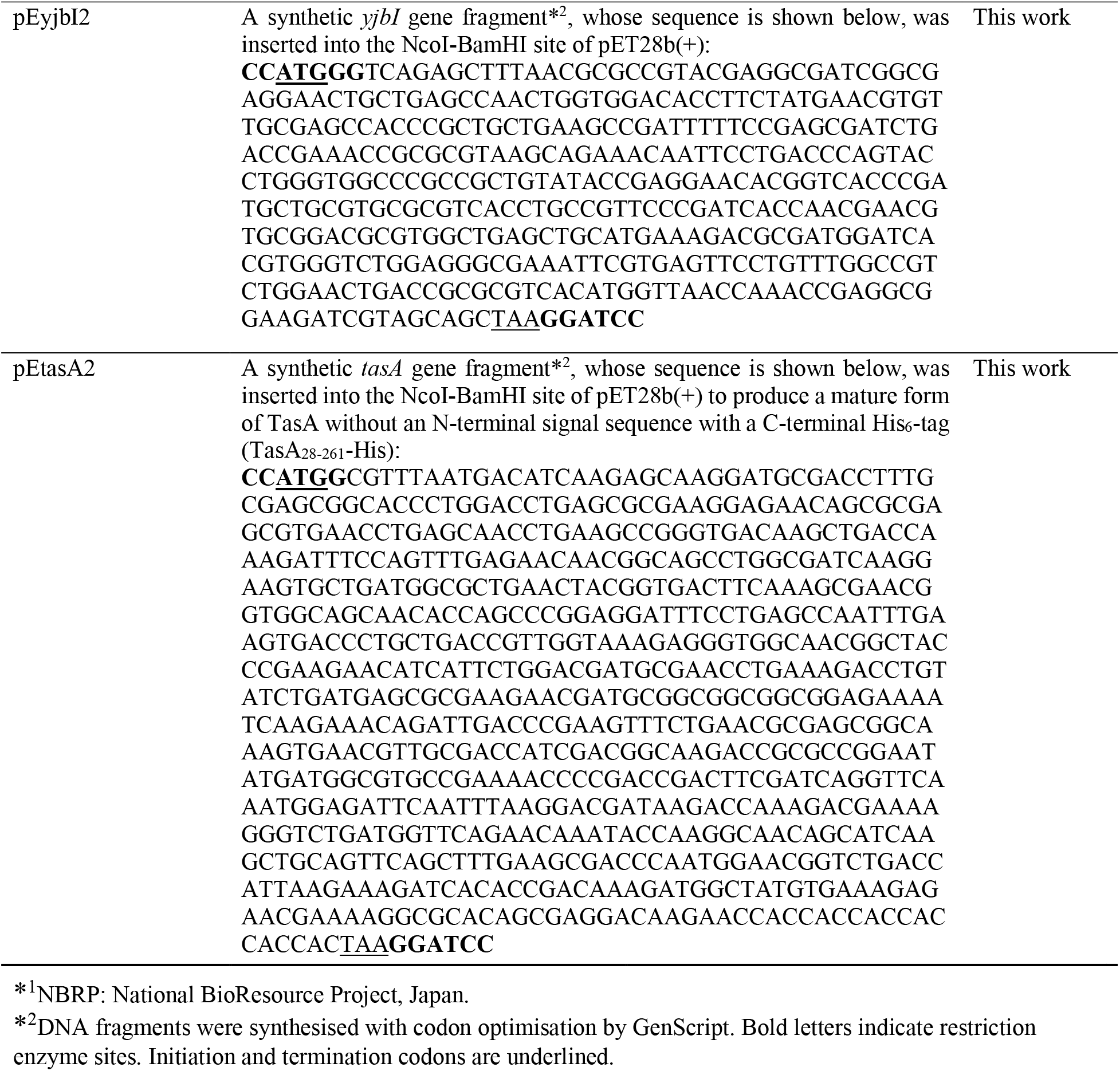
Strains and plasmids used in this study.

